# Biodistribution and inflammatory response to intracranial delivery of scintillating nanoparticles

**DOI:** 10.1101/609354

**Authors:** Máté Fischer, Amber Zimmerman, Eric Zhang, Joseph Kolis, Ashley Dickey, Mary K. Burdette, Mitra Afaghpour-Becklund, Praveen Chander, Stephen H. Foulger, Jonathan L. Brigman, Jason. P. Weick

## Abstract

Nanoparticle-based delivery systems have become a popular method for targeting tumors and impermeable tissue with drugs for treatment and imaging markers for biodetection. Nanomaterials are beneficial for medical treatment because they can be modified to have increased stability and carrying capacity, and their size and surface modifications allow them to reach otherwise impenetrable tissue. Localized or systemic injection can be used for delivery of nanoparticles; however, systemic injection without appropriate surface modifications is subjected to uptake by the mononuclear phagocytic system which clears particles from the circulation rapidly limiting their accumulation at target tissue and reducing efficacy. Here we demonstrate the biodistribution of Yttrium oxyorthosilicate nanoparticles doped with Cerium after localized injection to the cerebral cortex as well as the immune response at the site of injection over time.

## Introduction

Nanoparticles have become an increasingly popular tool for many applications ranging from diagnostics to drug delivery and targeted therapies (Soppimath 2001, Faraji 2009, Bahrami 2017). Much research has gone into the development of nanomaterials that are biocompatible and do not elicit a deleterious immune response. Despite this enormous promise of this emerging technology, issues with the delivery, targeting, toxicity and clearance of nanoparticle-based drug delivery systems leave many unresolved questions (Longmire 2008, Ray 2009). One strategy currently in development is to modify the surface chemistries of nanoparticles in such a way as to address these concerns, aiming to alter particles in ways that promote stability and minimize toxicity (Kango 2013).

In addition to modifying surface chemistries, researchers have also taken advantage of the fact that nanoparticles generated from different starting materials also display an enormous amount of variability in their biocompatibility and utility (Faraji 2009). Silicon-based nanoparticles have been especially extensively probed for potential medical applications due to their low inherent toxicity, especially when administered chronically (Murugadoss 2017). Wide band gap materials doped with rare earth ions are under intense research for their applications in medical imaging and have been used in the development of PET detectors for their scintillating properties (Loudyi 2007). More recently however, rare earth orthosilicates such as yttrium orthosilicate doped with cerium Y_2_SiO_5_:Ce (YSO:Ce) have been studied for their additional scintillating properties (Ricci 2008, Hamroun 2018). Scintillating nanoparticles provide a popular platform for biomedical applications because of their ability to produce light upon stimulation that can be detected through multiple imaging modalities. Scintillating materials such as YSO:Ce allow investigators to deliver specific spectra of light to targeted areas that can be utilized in applications such as optogenetic manipulations (Melcher 2000, Pignalosa 2012). YSO:Ce’s specific blue light emission spectrum, typically used in LEDs and phosphors, may provide a novel light delivery method to the brain inhibited by minimal light scatter (Krämer 2003, Gonzales-Ortega 2005).

Currently, in vivo optogenetic methods require the use of an implantable optical fiber into a target tissue expressing a genetically encoded bacterial opsin consisting of a channel protein capable of excitation upon stimulation with specific wavelengths (Cardin 2010). Scintillating nanoparticles could potentially be designed which emit a tunable spectra of visible light to replace currently used optical fiber-based approaches. Cell type specific expression of optically-activated constructs paired with light stimulation from localized scintillating nanoparticles provides a far less invasive platform for optogenetic experiments than current available methods of intracranial light delivery, such as surgical implantation of optical fibers. Previous work has demonstrated that only a median of 0.7% of systemically injected nanoparticles passively reached their target tissue, which limits the efficacy of these particles to barely appreciable levels (Wilhelm 2016). In addition to issues of ineffective passive distribution, the mononuclear phagocytic system also represents a significant barrier to the focal delivery of particles to target tissues (Wei 2018). Intracranial injection of nanoparticles to a target region would bypass the phagocytic system and allow for specific localization of particles to a brain regions of interest. Additionally, nanoparticles can be designed with surface modifications that tether them in close proximity to their sensors to provide more directed light stimulation in a cell-specific manner and increase their chemical stability.

Despite these promises, little is currently known regarding the biodistribution or toxicity of scintillating particles delivered via intracranial injection. In the current study, we examined the biodistribution of, and inflammatory response mounted against scintillating nanoparticles delivered into mouse cerebral cortex in order to determine the feasibility and efficacy of utilizing these particles for in vivo stimulation in future studies. C57BL/6J mice were given intracranial delivery of YSO-BSA-fluorescein (FITC) nanoparticles at differing concentrations targeting premotor cortex (M2) as it is a commonly targeted area in optogenetic experiments. Follow infusion, we examined the distribution and neuro inflammatory response was measured each at specific post-infusion time points.

## Methods

### Synthesis of Yttrium orthosilicate nanoparticles

Sub 100 nm silica cores were synthesized by the sol-gel process with 27 mmol (6.0 mL) of tetra ethyorthosilicate, 167 mmol (3 mL) of water, and 50 mmol (2 mL) of ammonia hydroxide (28% v/v) in 150 mL of ethanol at room temperature under vigorous stirring condition for 20 hrs. The particles were then washed with ethanol (2x) and water (1x). Silica (120 mg) was disperse in water (100 mL) with 1.98 mmol (760 mg) of yttrium nitrate hexahydrate and 15 μmol (6.5 mg) of cerium nitrate hexahydrate. Ammonia hydroxide (1 mL) was added drop wise into the solution and vigorously stirred for 16 hrs in room temperature. The waters were washed with water (3x) and dried at 125OC. The particles were annealed at 1000°C for 1 hr.

### Characterization of core-shell yttrium orthosilicate nanoparticles

Sub 100 nm particulates were dispersed in water on double-sided carbon tape and mounted onto a scanning electron microscope stub. The particles were plasma coated with platinum. A Hitachi 4800 scanning electron microscope was used to image the particles at 20 kV and 10 mA. Photoluminescence and excitation measurements were taken by a Jobin Yvon Fluorolog 3-222 spectrometer with a pump source of 360 nm. X-ray luminescence was measured with a Horiba JY synapse CCD camera, Horiba JY microHR monochromator, and a tungsten target Amptek mini x-ray source at 40 kV and 99 μA.

### Experimental subjects

Female C57BL/6J mice (The Jackson Laboratory, Bar Harbor, ME) were housed in a temperature- and humidity- controlled vivarium under a reverse 12 h light/dark cycle (lights off 0800 h). All experimental procedures were performed in accordance with the National Institutes of Health Guide for Care and Use of Laboratory Animals and were approved by the University of New Mexico Health Sciences Center Institutional Animal Care and Use Committee.

### Microinjection of Nanoparticles

At approximately 6 months of age, mice were anesthetized with isoflurane and fixed in a stereotaxic apparatus (1900 Stereotaxic Alignment System, David Kopf Instruments, Tujunga, CA) as previously described (Brigman & Josey 2015). A 33-gauge infusion cannula (Plastics One, Roanoke, VA) attached with polyurethane tubing to a Hamilton syringe (Hamilton, Reno NV) was directed at secondary motor cortex (M2. AP: 1.345, ML: +/− 0.6, DV: − 1.0). YSO-BSA-Fluorescein nanoparticles (0.5 μl) saline vehicle was infused over 5 min using a pump (GenieTouch, Kent Scientific, Torrington, CT), with the cannula left in place for an additional 2.5 min to allow full diffusion. Mice were injected with 3mg/ml RLPs in a saline solution and saline alone in contralateral hemispheres. On completion of the last infusion, mice were sutured and returned to their home cages.

### Immunohistochemistry

At 24, 72 or 216 hours (9 days) post-injection, mice were sacrificed by cervical dislocation, brains were extracted and placed in 4% paraformaldehyde solution for 24 hours. 50 μm coronal sections were cut with a vibratome (Classic 1000 model, Vibratome, Bannockburn, IL) adjacent to the site of injection. Slices were blocked and permeabilized for 1 hour on a Belly Dancer shaking platform (IBI Scientific) in a 500μL solution of 10% donkey serum, and 0.5% Triton X-100, brought to volume with 1x PBS. After one hour incubation at room temperature the blocking solution was removed and replaced with 500μL of primary antibody solution containing monoclonal mouse anti-GFAP (dilution: 1:500; Neuromab, USA) and polyclonal rabbit anti-IBA-1 (dilution: 1:250; Wako, Japan) diluted in 5% donkey serum and 0.25% Triton X-100 brought to volume in PBS. Sections were covered and incubated in a 4°C walk-in cold room overnight. The following day, the primary antibody solution was removed and slices were washed 3 times with 500μL PBS for 5 minutes each. Following the three washes, 500μL of secondary antibody solution containing donkey anti-mouse AlexaFluor 647 (dilution: 1:500; Abcam, USA) and donkey anti-rabbit AlexaFluor 550 (dilution: 1:500; Abcam, USA) diluted in 5% donkey serum and 0.25% Triton X-100 brought to volume in PBS was added to the sections at room temperature for 1 hour. After this incubation the secondary antibody solution was removed and slices were washed 3 times with 500 μL 1xPBS for 5 minutes each. DAPI counterstaining (1:5000 concentration in PBS) was then performed and slices were placed on a rocker for 10 minutes shielded from light. Finally, the slices were washed 3 times with 500μL PBS for 5 minutes each and mounted to glass microscope slides. Slices were covered with 100μL of Fluoromount-G (Thermo Fischer Scientific, USA) and then sealed with glass coverslips, making sure to remove any air bubbles. Slides were finally stored in a 4°C refrigerator for subsequent imaging.

### Confocal Microscopy

Following immunostaining and mounting, nanoparticle-tagged FITC and fluorescence-tagged immune cell populations were imaged using a Leica TCS SP8 Confocal Microscope (Leica Microsystems). DAPI, nanoparticle-conjugated FITC and secondary antibody-conjugated fluorophores were excited using a 405nm laser diode and a tunable pulse white light laser, respectively, with a 10x air objective and a 63x oil immersion objective and captured using hybrid spectral detectors (HyD). Z-stacks were then collected with a Z-range of 70μm. Z stacks of adjacent slices were ordered and merged to estimate the extent of the nanoparticle spread and compressed then into a two dimensional image using the maximal intensity projection algorithm in the Leica LASX software package. Maximum projection maps were exported as uncompressed TIFF files and transferred to ImageJ image analysis software.

### Analysis and Statistics

Optical sections of 25μm thickness were collected for the brains that were sectioned and stained for quantification of NP-FITC, IBA-1 and GFAP signals, respectively. After Z stacks were compressed, ImageJ software was used to draw regions of interest (ROIs) surrounding the entire extent of the M2 region with the help of the Allen Brain Atlas for the given appropriate A/P range. Light signal from photon scatter around the edges of tissue, tears in the tissue as wells as vasculature were excluded from analysis. Mean fluorescence intensities were calculated over the ROIs, and a background measurement taken from cortical regions outside M2 was subtracted from these values to normalize for the autofluorescent and nonspecific signal present in the sample. A ratio was then calculated for signal intensity in the NP vs. vehicle treated hemispheres and mean values with SEM were reported for each time point. ImageJ was scaled such that the pixel to mm ratio was defined for each image and using this scale, the area of the extent of NP spread was calculated by manually tracing the outlines of the FITC signal adjacent to the site of injection. Two-tailed, unpaired t-tests were conducted to compare replicate MFI values across time points.

## Results

To determine the extent of spread of injected nanoparticles, animals that received the particles were sacrificed 24 hours, 48 hours and 9 days post injection and the FITC fluorophore conjugated to the nanoparticles was imaged across the extent of the M2 region. Confocal imaging of the region surrounding the injection site revealed reliably detectable FITC in non-perfused animals (Figure 3A, D), but no detectable signal in animals that were perfused with 4% PFA upon sacrifice (data not shown). In the non-perfused groups, 50μm slices were immunostained for the astrocytic marker glial fibrillary acidic protein (GFAP) as well as the microglial marker allograft inflammatory marker 1 (IBA-1) for subsequent analyses (Figure 3B, E). Due to light scatter in the sample along edges and tears in the tissue, as well as from vasculature giving off fluorescent signal in addition to our sample, care was taken to exclude these confounds from analysis (Figure 3C, D). With this in mind, we set out to quantify the area and intensity of the spread of NPs as well as the presence of inflammatory cells in the M2 region.

**Figure 1.**
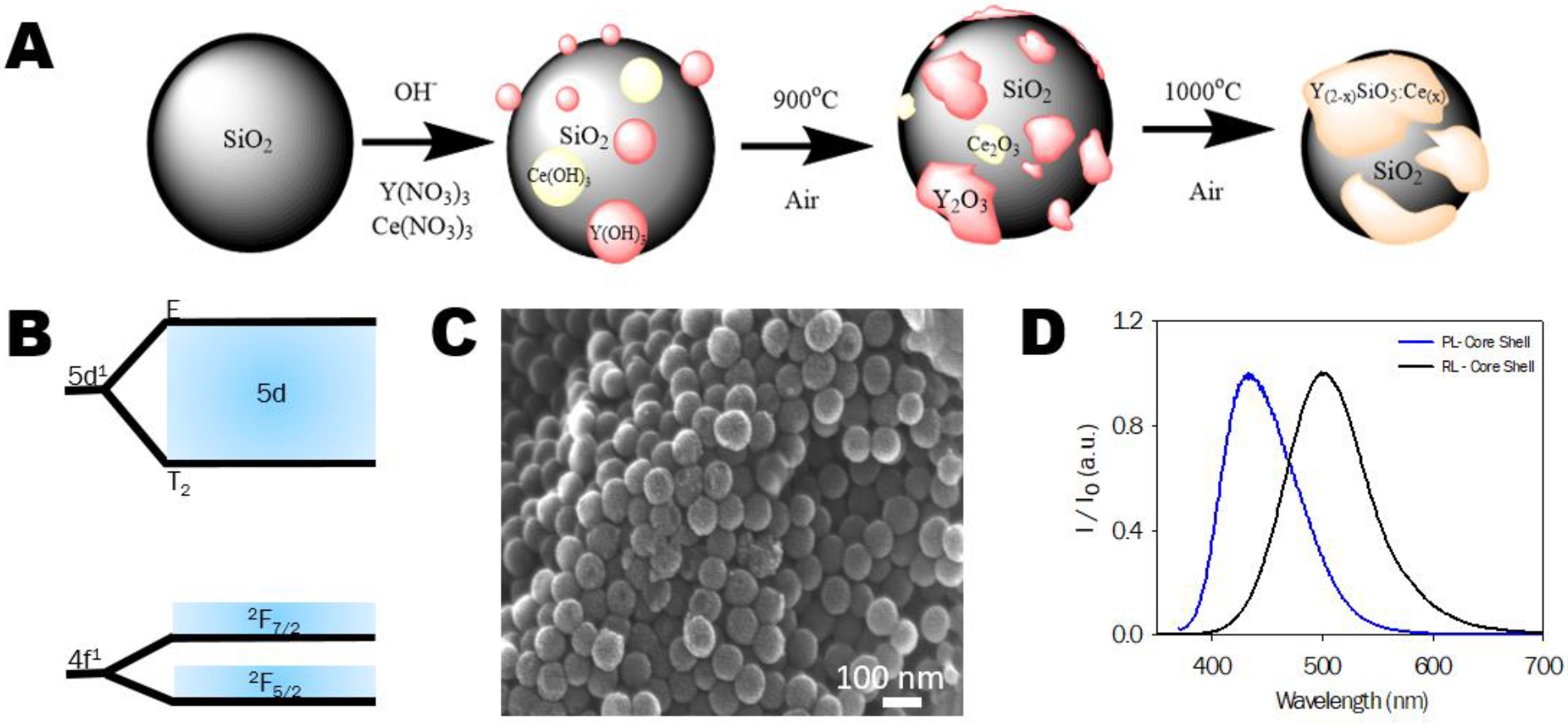
A) Schematic of the development of YSO:Ce using a core-shell method. B) Relaxation of cerium from it higher energy 5d state to its respective ground state of ^2^F_7/2_ and ^2^F_5/2_ in the Ce 1 site. C) SEM of NPs synthesized by the core-shell method. D) Photoluminescence and radioluminescence of nano core shell particulates irradiated by 360 nm and a broadband 40 KeV tungsten x-ray source, respectively.

**Figure 2.**
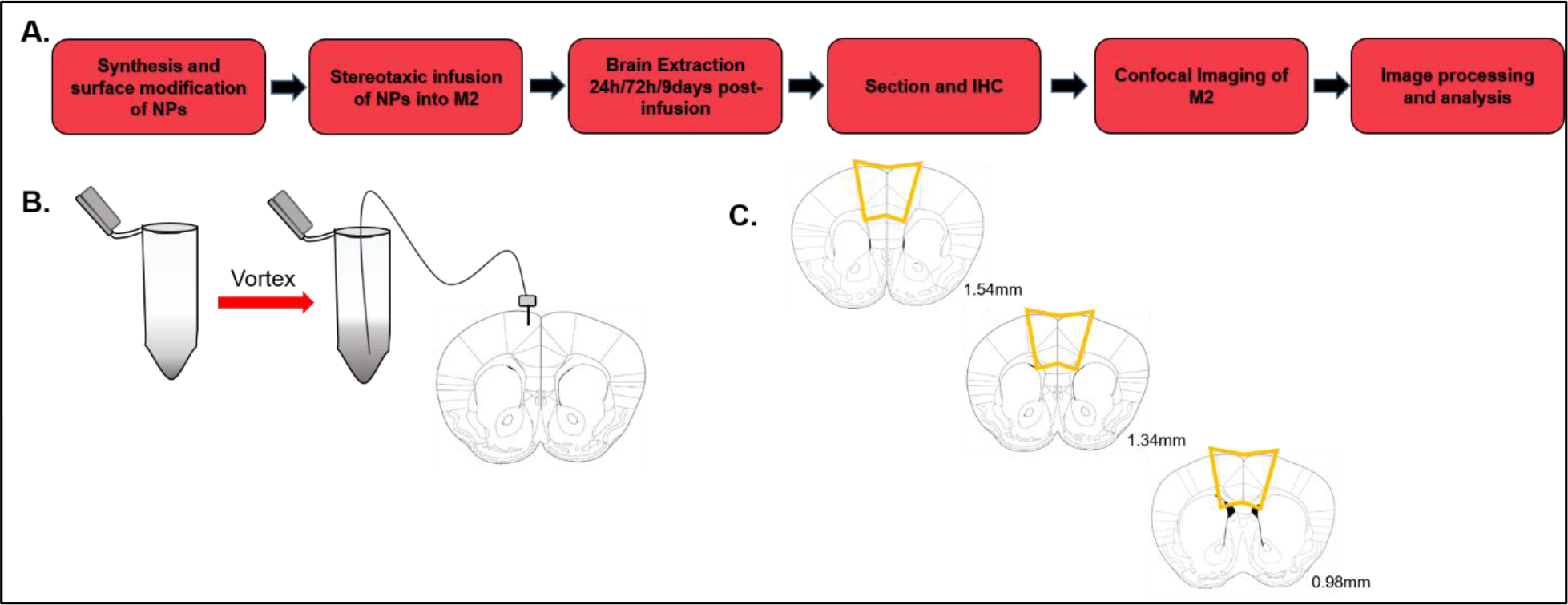
A) A. Workflow describing synthesis and infusion of nanoparticles across 3 time points to analyze biodistribution from 1 to 9 days. B) B. Intracranial nanoparticle injection location (AP: +1.34; ML: +/− 0.60; DV: −1.00). Nanoparticles aggregate and should be vortexed thoroughly (10 seconds) immediately before injection to maximize particle distribution. C) C. Typical range of nanoparticle spread throughout cortex. Orange polygon represents area of quantification for confocal microscopy.

**Figure 3.**
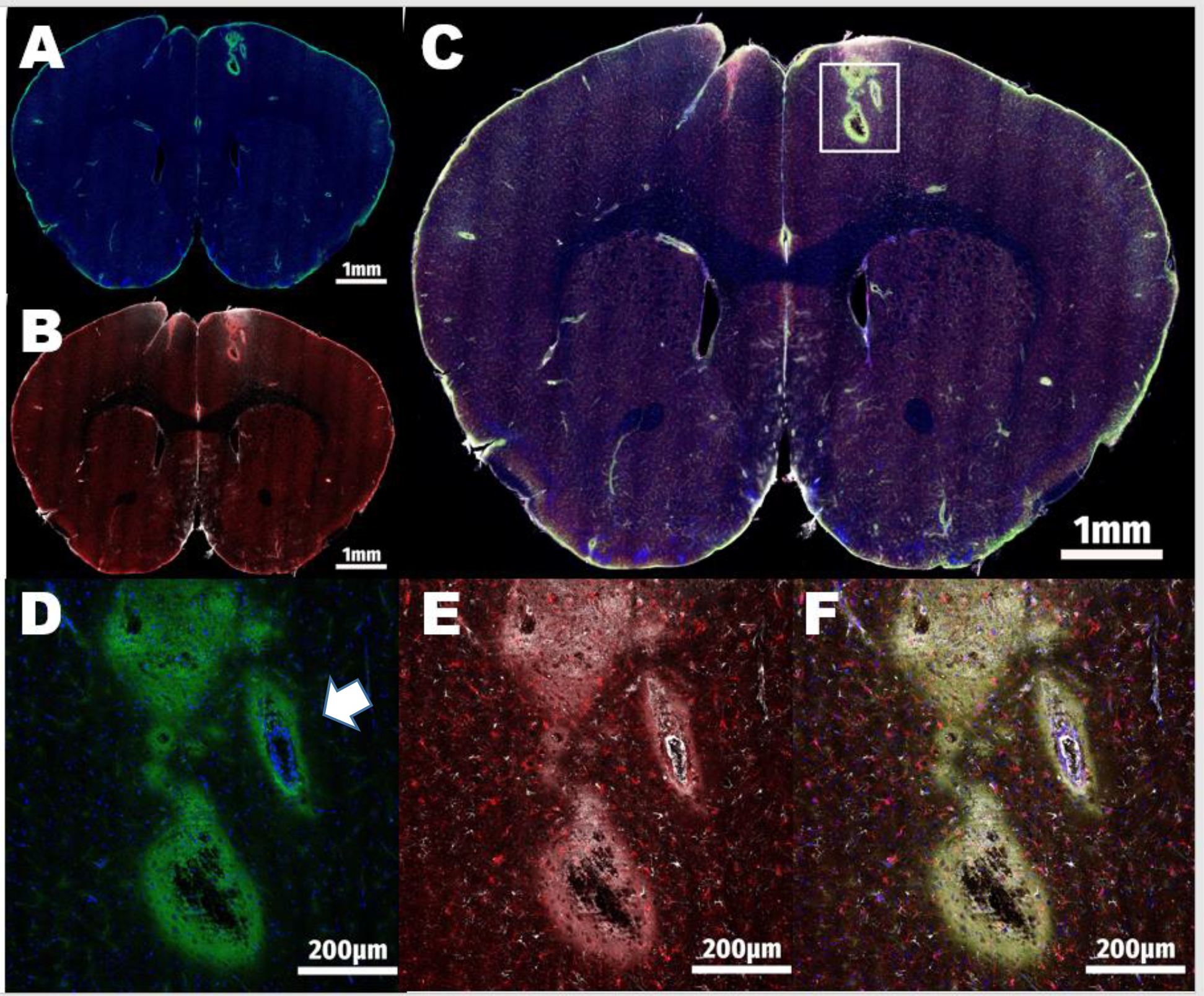
A) Full 40x tile scan of NP-injected mouse brain showing NP-FITC in green and nuclear DAPI staining in blue. The green signal around the edges of the tissue due to light scatter was excluded from analysis B) Same mouse brain showing IBA-1 (red) staining as well as GFAP (white) C) Merge of NPs (green), DAPI (blue), IBA-1 (red) and GFAP (white) staining as well as highlighting area of inset shown in D-F D) NP and DAPI staining show in 63x tiled insert. White arrow is indicating a blood vessel excluded from analysis E) Regularly distributed IBA-1-expressing (red) microglia are clearly visible at higher magnification as well as GFAP-expressing (white) astrocytes F)4 channel merge demonstrates the area of extent of RLP spread as well as the presence of neural immune cells

To accomplish this quantification, Z stacks were collected across the extent of the M2- containing brain sections and 3D interpretations were reconstructed with Leica LASX software (Figure 4A). Based on this 3D reconstruction it was determined that the injected NPs spread to fill an area of approximately 0.091mm3 in the 24hr group, 0.097mm3 in the 48hr group and just 0.0101mm3 in the long term group (Figure 4B). The drop in signal intensity adjacent to the injection site between the 72 hour and 9 day time points could potentially indicate either an active clearance of the particles to the lymphatic system, or a passive diffusion of the particles through the M2 region away from the injury site between those two time points. To address this difference in mechanisms, Z stacks were then flattened according to a maximal intensity projection algorithm and ImageJ software was utilized to measure mean fluorescence intensity (MFI) for the NP-FITC signaling across the entire extent of the M2 region from both the NP injected (ipsilateral) and vehicle-injected (contralateral) sides. These fluorescence measurements were normalized to a cortical background fluorescence level outside the M2 region to control for nonspecific signal, and are shown as a ratio of the intensity of the FITC signal in hemisphere ipsilateral to the injection over the contralateral side (Figure 4C). Interestingly, while the size of the relatively bright NP-FITC signal adjacent to the injection site diminished notably by the 9 day time point, no difference was established in the overall mean fluorescence intensity across M2 in the same time frame, indicating the likely persistence of NPs in the M2 region, likely due to passive diffusion from the site rather than clearance from the tissue overall.

**Figure 4.**
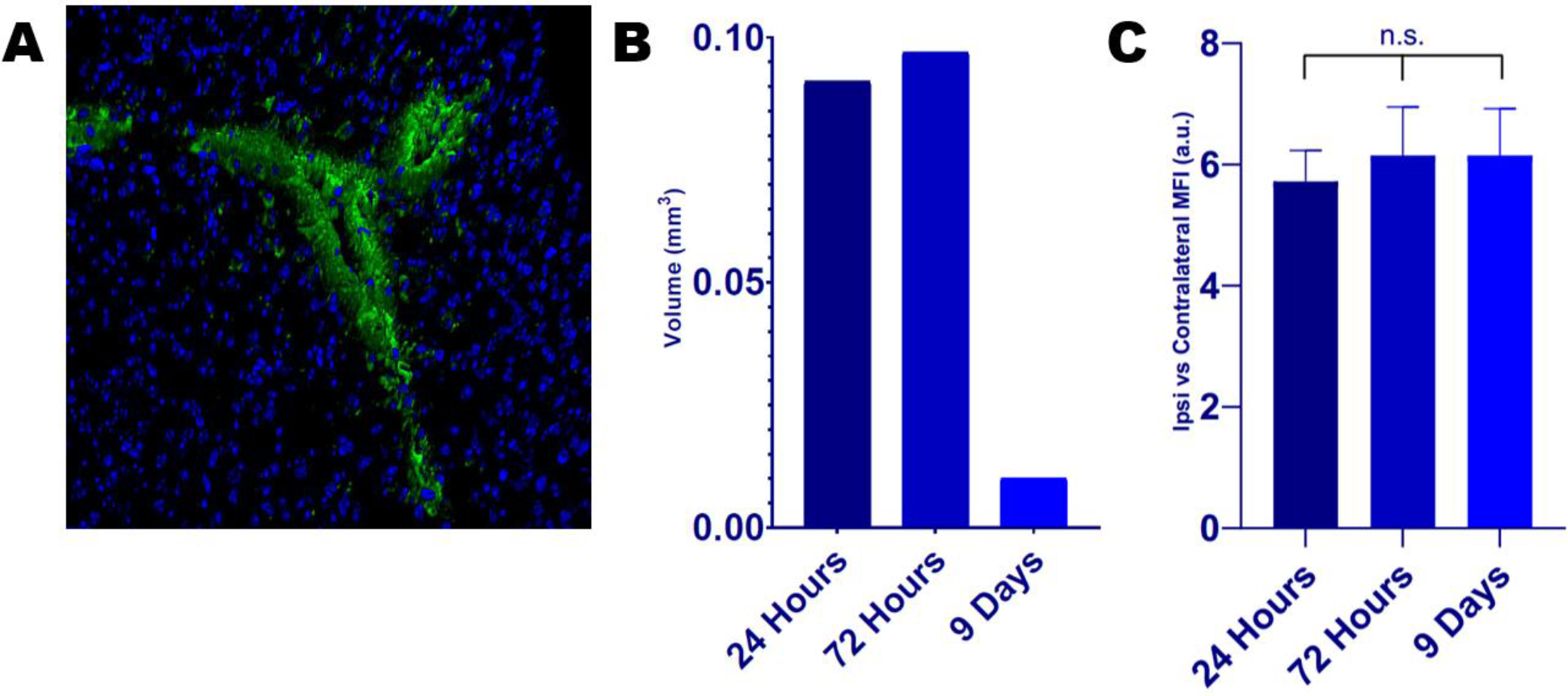
A) 3D render of an injection site, representative of the analysis used to approximate the spread of NPs from the M2 injection site B) Time course of the area of extent of bright NP signal adjacent to the injection site shows no change across the first 72 hours, but a sharp decline at 9 days, indicating diffusion or clearance from the injury site in that time C) Interestingly, the ratio of NP fluorescence of the ipsi- to contralateral side remained unchanged over the 9 days assaye/given-names>, suggesting the persistent presence of NPs despite the diffusion from the injection site

As previously noted, brain sections were additionally stained with antibodies against astrocyte marker GFAP as well as microglial marker IBA-1 near the injection site to determine whether or not the brain immune system mounted an inflammatory response to the presence of the particles. Representative images for each the IBA-1 (Figure 5A) and the GFAP (Figure 5B) staining, respectively, show that both glial cell types were readily detectable in the mouse gray matter following NP injection. Maximal projection maps were again generated for the region of interest and mean fluorescence intensities over the M2 region were determined for each time point. According to these quantifications, IBA-1 intensity was significantly transiently increased between 24 hours post-injection (p=0.0375) and then returns to baseline levels by the latter time point (p=0.0475) (Figure 5C). Presence of GFAP^+^astrocytes also showed temporal regulation following NP injection, but in this case demonstrated a maximal response 24 hours following the particle delivery, only to return to baseline levels by the 72 hour mark (p<0.001) and stay low at the 9 day time point (p<0.001) (Figure 5D).

**Figure 5.**
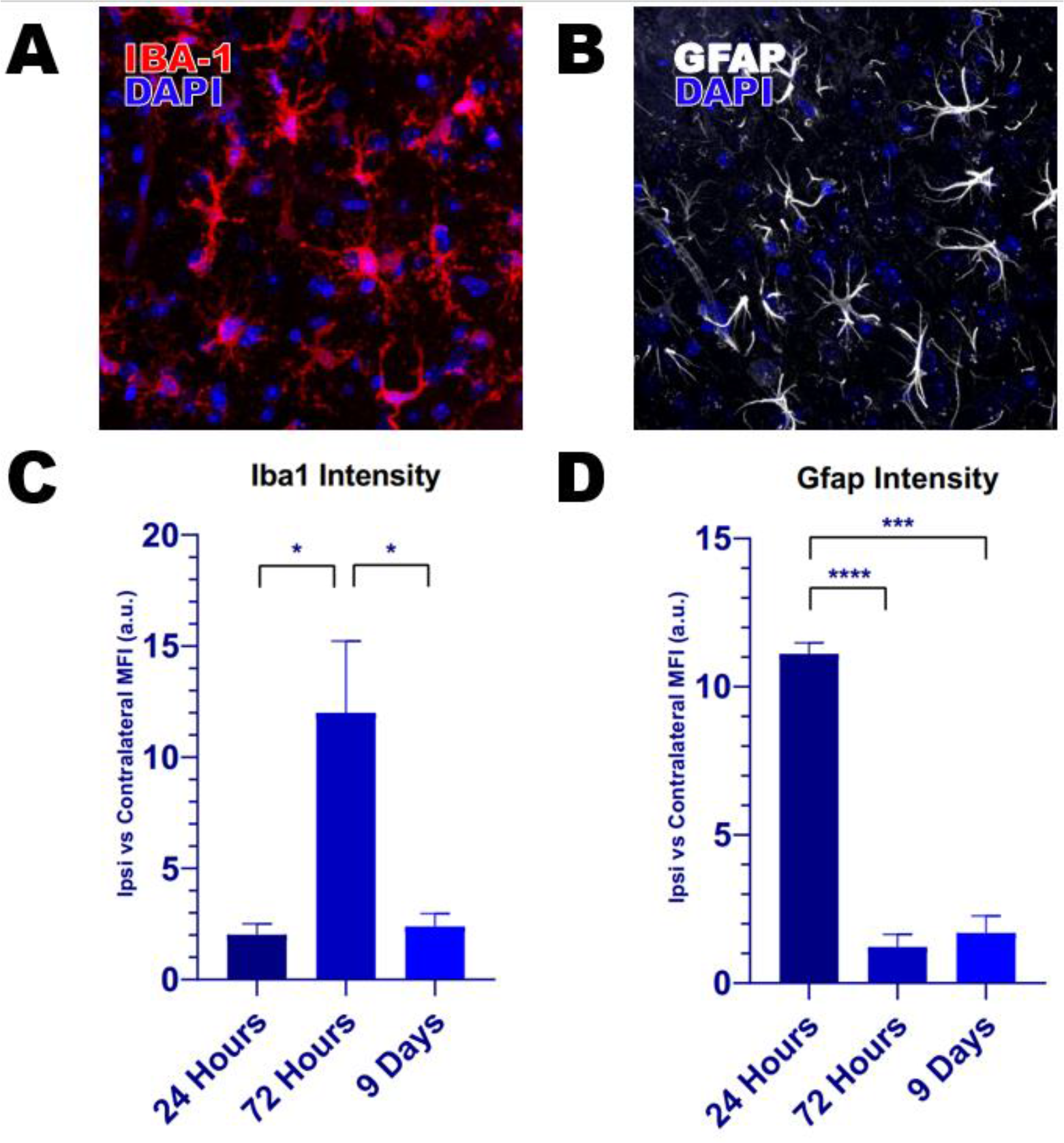
A) Representative image of IBA-1 staining for microglia B) Representative image of GFAP staining for astrocytes C) Time course of IBA-1 intensity shows an elevation in microglial staining across the entire M2 region at 72 hours post-injection, but not 24 hours or 9 days D) Time course of GFAP intensity shows an elevation in astrocytic staining across the entire M2 region 24 hours post-injection, but not at later time points

While our data do suggest that an immune response is mounted against the particles that is greater in magnitude than that of our control saline injection, it seems that after 9 days in the brain, any inflammatory response has been normalized and the particles not only diffuse passively from the site of injection, but remain in the brain to an extent indistinguishable from merely 24 hours after the NPs are delivered. These findings taken together suggest a potential utility for injectable YSO:Ce nanoparticle platform in the delivery of light or any surface modified residue to brain tissue with a minimal, transient induction of neuroinflammation that may be superior to long-term surgical interventions currently required for optogenetic, electrode arrays and other applications for probing murine neural function and networks.

## Discussion

From drug and gene delivery to imaging to optical stimulation, nanoparticles could serve as a platform technology for a new wave of medical and research initiatives (Singh 2011). Inherent properties of the particles such as photon emission by scintillation as well as diverse modifications to surface chemistry can be readily achieved, making the biological compatibility of such particles the next major hurdle for most preclinical and clinical research (Baizar 2011, Kulkarni & Feng 2011).

Some work has been done concerning the diffusion dynamics of novel nanoparticles, but wide variability between reports in NP size, composition and surface chemistry yield a wide heterogeneity of results (Neugart 2007, Yu 2011, Grabowski & Mukhopadhyay 2014). Additionally, material science groups synthesizing nanoparticles for biomedical applications frequently model dispersion dynamics *in silico* or in polymers, gels and fluids, but relatively few reports published to date include *in vitro* or *in vivo* diffusion data (Song 2009, Chauhan 2011, Nance 2012).

In this study, we demonstrate that LSO:Ce NPs persist at statistically unaltered levels through the 9-day post-injection time point, but some time after 72 hours the particles began to diffuse away from the site of the injection, while remaining detectable in the M2 cortical region. This suggests that the particles discussed in this report yield promising preliminary results for the diffusion potential of the particles though gray matter to target larger cortical or deeper brain regions over longer time points. In line with previous reports, the sub-200nm size of the NPs utilized should readily diffuse through the brain extracellular space (Nance 2012). The diffusion of the NPs paired with the finding that they are not being cleared from the brain begs the question of to what degree the local microenvironment and relevant inflammatory factors will respond deleteriously the injected particles.

Perhaps most interesting among our findings concerning the neuroimmune response to local injection of nanoparticles was the temporal regulation of GFAP and IBA-1-positive cell populations. The response of astrocytes to a stab wound in the cerebral cortex of rodents has long been appreciated (Mathewson 1985, Eclander 1990, Hatten 1991). While our results do not preclude an effect prior to 24 hours, and indeed, astroglial response to focal mechanical trauma can be detected within an hour of injury according to some reports, we do observed that any increase in the presence of GFAP^+^astrocytes had subsided by the 72 hour and 9 day time points (Eng 1994). Acute stab injury alone has been shown to only minimally induce astroglial presence in grey matter, however the inclusion of foreign material such a nitrocellulose, as reported by Balasingam *et al.*, can induce robust recruitment and proliferation of astrocytes (1996).

Once localized to the lesion, astrocytes are known to remodel the injury site itself in a number of ways, including alterations to blood-brain barrier permeability, glial scar formation as well as a number of changes to the tissue and extracellular matrix (ECM) structure and composition (Johnson 2015, Burda 2016). These ECM alterations, primarily thought to facilitate clearance of cellular debris could underlie the diffusion of NPs away from the injection site as well as the subsequent recruitment of microglia, levels of which were shown to be elevated at the later 9 day time point, but not while GFAP levels were raised 72 hours post-injection.

In addition to surface modifications designed to facilitate biocompatibility and distribution of locally delivered nanoparticles, inherent properties of the materials at the core of the NPs also hold potential value to investigators. In this instance, LSO:Ce-based nanoparticles were utilized for their potential to emit blue photons upon excitation with low levels of ultraviolet (UV) or x-ray stimulation, potentially useful for optogenetic stimulation of cohorts of neurons of interest. To this end, Bartley *et al.* observed that light emitted from LSO:Ce nanoparticles in response to UV stimulation caused an increase in the frequency of sEPSCs onto CA1 pyramidal cells expressing the light sensitive opsin channelrhodopsin-2 (ChR2), commonly used for optogenetic experiments (Boyden 2005, Bartley 2019). However, Bartley *et al.* were not able to detect firing of action potentials or large light-evoked EPSCs, consistent with the relatively small photocurrents observed. Furthermore, Bartley and colleagues did observe small changes in synaptic function including a small reduction in the frequency of excitatory postsynaptic currents (EPSCs) and an increase in paired pulse ratio of evoked excitatory transmission, indicating an effect of the particles on presynaptic release probability (Dobrunz & Stevens 1997, Crabtree 2017), suggesting that this could be the underlying mechanism for the minor effects on synaptic function with application of the LSO:Ce NPs. It is important to note that NPs for these experiments were applied to the surface of acute slice preparations, with estimated distance from recorded ChR2-expressing cells of 100 microns or more. Our results support the notion that direct injection of particles could be used prior to slice preparation to directly test for greater ChR2 activation due to closer proximity and reduced light attenuation from scattering (Yona 2016). Taken together, this report finds that the nanoparticles described here readily diffuse away from the site of injection in grey matter, while persisting in the brain up 9 days post-injection and causing a temporally regulated immune response that is no longer detectable in the injected cortex 9 days after their delivery. Particles with these properties should be highly desirable for applications of drug delivery to target brain regions or light emission for use in optogenetic studies.

## Acknowledgements

This project was supported by an NSF EPSCoR track II award (NSF1632881)

